# Enhancing the imaging rate of high-speed atomic force microscopy using a combination of multiple techniques

**DOI:** 10.64898/2025.12.22.695957

**Authors:** Shingo Fukuda, Akihiro Otomo, Ryota Iino, Toshio Ando

## Abstract

Protein molecules, while functioning, undergo dynamic conformational changes and interactions with their partners, which can be directly observed by high-speed atomic force microscopy (HS-AFM). However, its imaging rate of approximately 10 frames per second (fps) has remained unchanged since its inception, despite the increasing demand for observing dynamic biomolecular processes that occur too fast to be captured at this time resolution. Here we report a combination of techniques that significantly enhances the imaging rate without disturbing the sample. HS-AFM imaging of the stator ring of F_1_-ATPase visualised two intermediate states, which could not be resolved with the previous HS-AFM system. The high-resolution molecular movies demonstrate how the catalytic subunits cooperate during the chemomechanical cycle.

---

Structural biology is currently experiencing a period of significant transformation. The focus is shifting from static to dynamic structures^1^. The development of high-speed atomic force microscopy (HS-AFM), which has been achieved via substantial improvement on traditional AFM, has enabled the imaging of biological macromolecules in dynamic action in approximately 100 milliseconds^2,3^. Since its establishment in 2008^4^, HS-AFM has been used to visualise various dynamic molecular processes of proteins and nucleic acids, thereby accelerating our understanding of how they function^5–10^. Similarly, cryogenic electron microscopy (cryo-EM) has recently evolved into time-resolved cryo-EM (trEM) with a temporal resolution of 5−10 ms^11–16^, which surpasses that of HS-AFM. While the use of trEM has been demonstrated for only a limited number of targets to date, it will be used extensively for a wide range of targets in the future.

However, it should be noted that these dynamic microscopy techniques complement each other in terms of their respective strengths and weaknesses. For example, while HS-AFM visualises topographic structures, trEM can reveal three-dimensional structures with atomic resolution. HS-AFM can track single molecules over extended periods, but trEM is currently limited to acquiring time-evolving structures at a maximum of 4−5 time points, due to the time-consuming nature of structural determination. In addition, aligning the structures obtained by trEM along a time axis is challenging because multiple structures appear in the image ensemble at each time point. Furthermore, trEM presents challenges in the context of proteins exhibiting symmetry. For example, it is unable to detect 120° rotational conformational changes in F_1_-ATPase with threefold rotational symmetry. The limited speed performance of HS-AFM significantly restricts the range of targets to which it can be applied, and the use of HS-AFM data of a sample for understanding the time evolution of trEM structures of the same macromolecule is hampered.

Despite recent efforts to develop faster scanners^17,18^ and amplitude detectors^19,20^, the speed performance of HS-AFM has not significantly improved since its establishment in 2008^4^. This is because the cantilever resonant frequency (*f*_c_) is the main factor limiting the imaging rate^2^. Nevertheless, it is very challenging to increase *f*_c_ significantly without stiffening the cantilever. Consequently, even when the response speed of all components is improved, the imaging rate can only increase by a factor of 2−3 (Ref. 2). Therefore, in addition to efforts to increase the response speed of cantilevers and scanners, it is imperative to find alternative solutions that can effectively increase the imaging rate of the total system.

In this study, we developed an instrument that combines several techniques, with the aim of bringing the time resolution of HS-AFM closer to that of trEM. It is important to note that achieving a high level of speed performance cannot be accomplished by a single technique. It is necessary to combine several techniques, even when each of these has no remarkable effect. The primary consideration in this development, in addition to enhancing the response speed of mechanical components, was that the sample’s edge regions with a steep uphill gradient along the X-direction are subjected to substantial force from the cantilever tip during rapid scanning of the sample stage in the X-direction. The integrated techniques employed mitigate this force. Consequently, fragile actin filaments, which could previously only be imaged at 3−7 frames per second (fps), can now be imaged at 33−70 fps, depending on the orientation of the actin filaments relative to the X-direction. The stator ring (α_3_β_3_ subcomplex) of a rotary motor F_1_-ATPase with a steep slope at its edge can now be imaged at 40 fps, revealing intermediate states that could not be resolved by the original instrument.

## RESULTS

### Shorter cantilevers

The resonant frequency (*f*_c_) of the cantilever affects how quickly it responds and updates its oscillation amplitude. This, in turn, has a direct impact on the rate at which the feedback signal is transmitted to the piezo driver for the Z-scanner. Therefore, the speed performance of HS-AFM^2^ is primarily determined by *f*_c_. The highest *f*_c_ in water to date has been achieved using short silicon nitride cantilevers (BL-AC7DS-KU2, Olympus, Japan; discontinued), which are 6–7 µm long, 2 µm wide and 90–100 nm thick. They are characterised by *f*_c_ = 2.4–3.6 MHz in air, *f*_c_ = 0.8–1.2 MHz in water, a spring constant (*k*_c_) of 0.15–0.2 N/m and a quality factor (*Q*_c_) of 1.5–2.0 in water^21^.

Although it is difficult to produce shorter cantilevers with a small spring constant, we obtained them by milling conventional silicon nitride cantilevers with a thickness of 130 nm using a focused ion beam (FIB) in collaboration with a manufacturer. The produced cantilevers are spoon-shaped, with a spoon head measuring 1 × 1 µm² and a handle with a length of 3.5 µm and a width of 400 nm (see Fig. 1a). Only the backside is coated with gold to achieve efficient photothermal excitation^22,23^. The resonant frequency measured in water is ∼2.0 MHz (Fig. 1b), which is close to one-third of the theoretically estimated value in air^24^ (6.3 MHz). They have *k*_c_ = 0.25 N/m and *Q*_c_ = 2.0 in water. Due to the small spoon head, the reflected laser power is ∼20% lower than that of conventional short cantilevers. Nevertheless, the optical beam deflection (OBD) sensitivity is 0.03 V/nm, approximately 70% of that of conventional short cantilevers. This relatively high sensitivity is due to the use of the quotient signal (V₁ – V₂)/(V₁ + V₂), where V₁ and V₂ represent the voltage signals from the two segments of the position-sensitive photodiode sensor. Shorter cantilevers can enhance the imaging rate by a factor of ∼2 (Ref. 2).

**Fig. 1.**
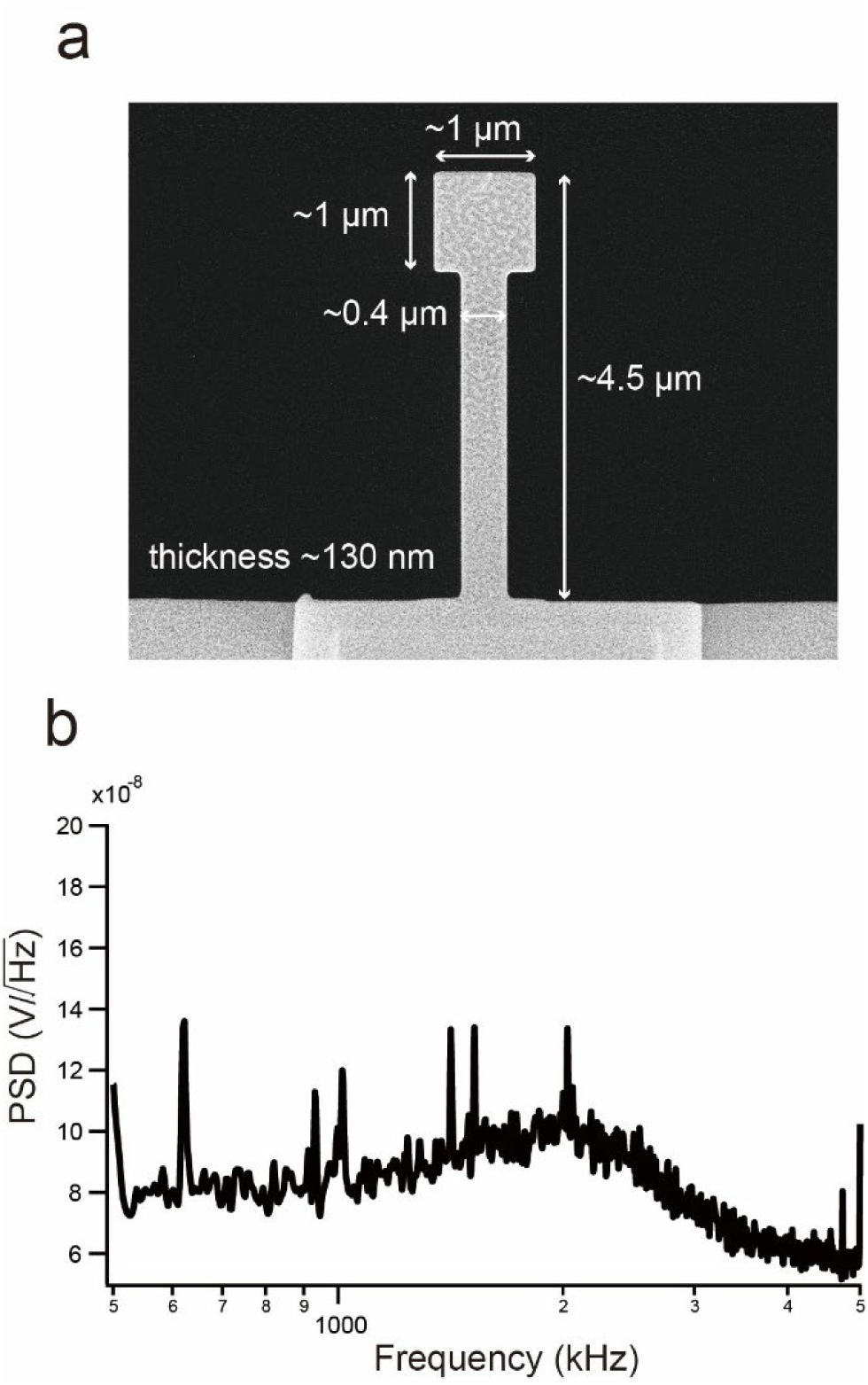
A spoon shaped cantilever. **a**, A scanning electron microscopy image of a spoon shaped cantilever. **b,** A frequency spectrum of thermal fluctuations of the spoon shaped cantilever.

### Reducing tip−sample contact force

In HS-AFM imaging, the cantilever tip is subjected to forces from the sample in both the X and Z directions, particularly in regions of the sample where the slope in the X-direction is steep. The cantilevers commonly used in HS-AFM (6−10 μm in length) are only 2–5 times longer than the 2–3 μm tip. The spoon-shaped cantilevers mentioned above are only 1.5–2.3 times longer. Therefore, the torque exerted on the cantilever by the X force is not considerably smaller than that exerted by the Z force. When the sample stage is being scanned forwards in the X direction, both torques act in the same direction. However, when scanning in the opposite direction, the two torques are exerted in opposite directions (see Fig. 2). Consequently, the cantilever does not deflect much during backward scanning, even when large X- and Z-forces are exerted. This results in insufficient or even opposite Z-scanning of the sample stage by feedback control. Therefore, if the sample is damaged during imaging, it is damaged predominantly during backward scanning, as demonstrated in our previous study^25^. An additional problem that should be addressed is the feedback gain. The sample surface corrugation exhibits various spatial frequencies in the X direction. However, the feedback gain remains constant when scanning over low-and high-frequency regions.

**Fig. 2.**
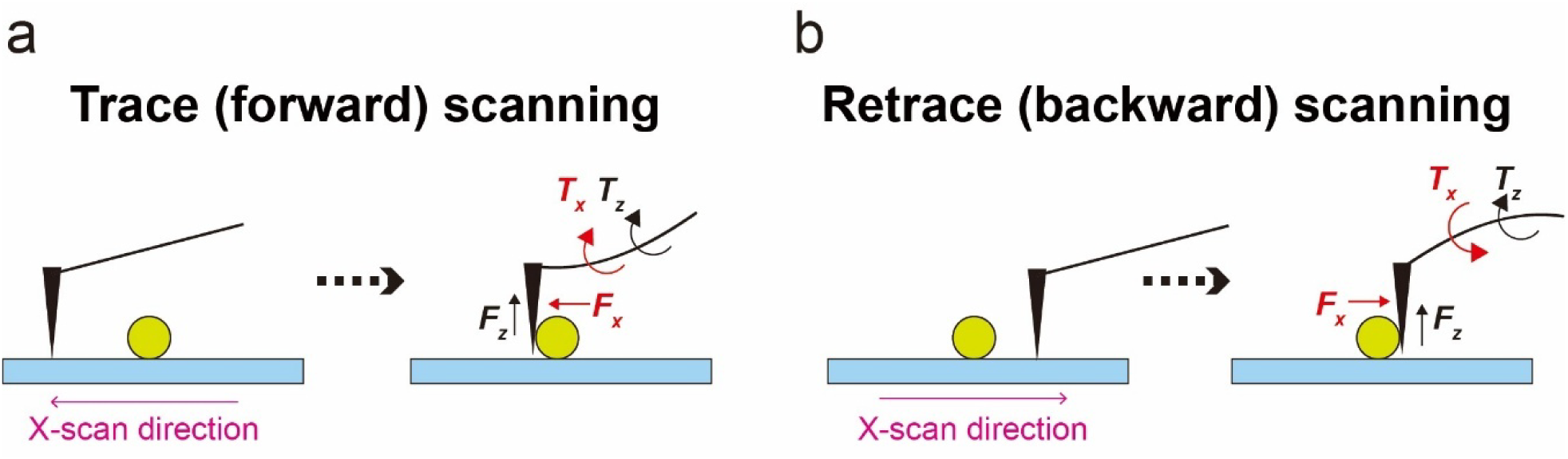
Schematic diagram of X-scanning in HS-AFM. **a, b**, Forward scanning during trace imaging and backward scanning during retrace imaging. The directions of torques generated on a cantilever by X- and Z-forces exerted from the sample to the tip are the same during trace imaging (**a**) but opposite during retrace imaging (**b**).

To address these issues, two strategies are required: the elimination of retrace imaging during backward scanning and the adjustment of the feedback gain according to the spatial frequency of the sample in the X direction. To address the first issue, we recently used ‘only trace imaging’ (OTI) mode^25^, whereby images were only captured during forward scanning. During backward scanning, the tip was lifted from the sample surface, and the scan velocity was increased. However, this approach was only moderately effective, increasing the highest possible imaging rate by no more than 2.5 times. The second strategy involved the use of a feedback controller capable of automatic gain tuning (AGT). This had no effect when it was previously tested, although it is effective in preventing complete tip detachment from the sample surface (’parachuting’) during scanning in descending regions of the sample^26^. The lack of effect of the AGT at ascending regions may be due to insufficient cantilever deflection during backward scanning. The AFM instrument interprets this insufficient deflection as low-spatial-frequency regions being scanned. Therefore, the combination of the OTI mode with the feedback controller with AGT should be more effective.

### Combining OTI mode with ATG

There are several methods that can be used for AGT in feedback control. We employed the simplest method, which had been designed previously^26^. A threshold level (*θ*_L_) is set below the set-point amplitude (*A*_s_). When the measured amplitude (*A*_m_) is smaller than the threshold level, the feedback gain (*g*_L_) is increased (see Fig. 3a). This condition (A_m_ < *θ*_L_) corresponds to a situation in which the corrugation of the sample surface in the current scan area has a high spatial frequency. To evaluate the sample protection effect of this method, we imaged fragile actin filaments aligned along the Y-axis or largely slanted from the X-axis for 10 seconds. In these orientations, the spatial frequency of the actin filaments is high in the X direction. Imaging was repeated 20−40 times under each condition to estimate the non-breaking probability (*P*_NB_)^25^. The spoon shaped cantilevers were oscillated photothermally using a laser (Supplementary Fig. 1). The free oscillation amplitude (*A*_0_) was set to 2 nm. The imaging was carried out in OTI mode. *P*_NB_ was obtained at different imaging rates, using the AGT feedback controller under the conditions of *A*_s_/*A*_0_ = 0.8, *θ*_L_/*A*_0_ = 0.6 and *g*_L_ = 2 (i.e. two-times higher gain than the gain used in the conventional feedback scheme). The results are shown in Fig. 3b (*P*_NB_ obtained under different conditions are given in Supplementary Fig. 2). The imaging rates giving *P*_NB_ = 0.8 were 24 (without AGT) and 33 fps (with AGT). Therefore, the imaging rate was extended by a factor of 1.4, without deterioration in the image quality (Fig. 3c). Note that the 24 fps without ATG is a result of the use of OTI mode, spoon-shaped cantilevers and a new Z-scanner with a resonant frequency of 350 kHz (it is 100 kHz in the original HS-AFM instrument). The imaging rate giving *P*_NB_ = 0.8 could be enhanced up to 50 fps by increasing *g*_L_, but the image quality was lowered (Supplementary Fig.3).

**Fig. 3.**
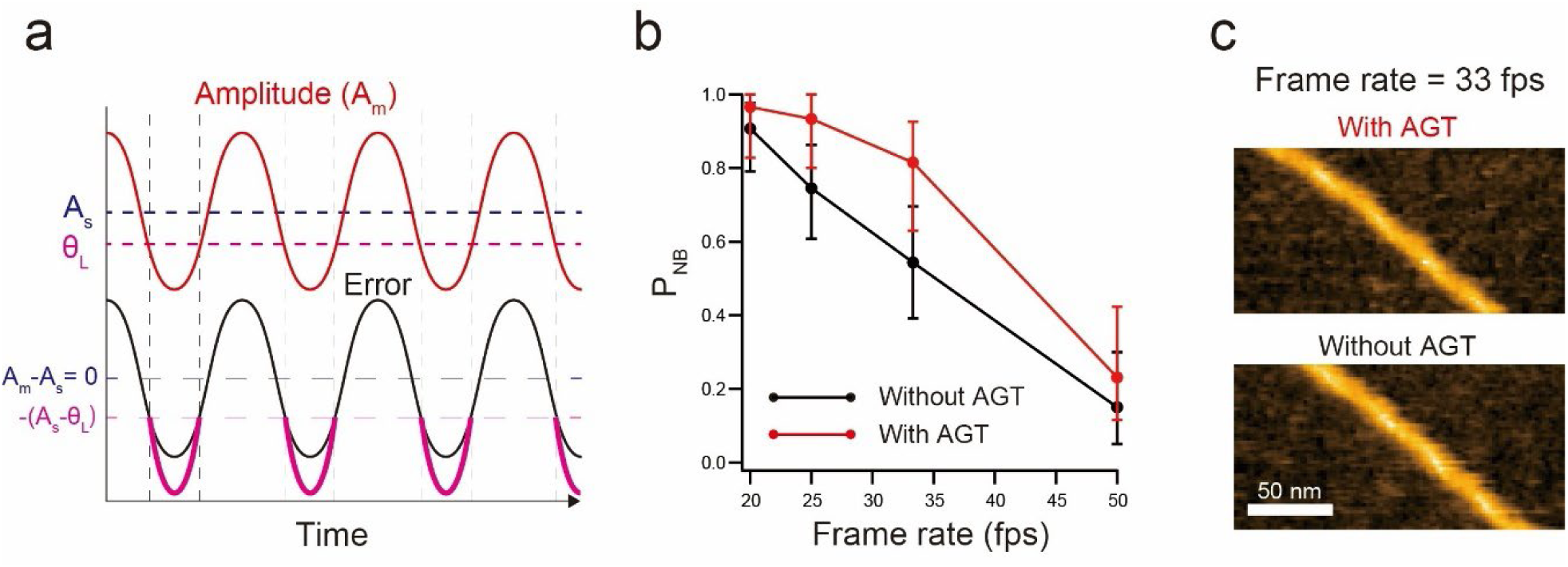
AGT feedback control. **a**, Schematic diagram for the AGT feedback control. **b,** Relationships of P_NB_ vs frame rate measured by imaging of actin filaments largely slanted from the X-axis unde feedback control with and without AGT. The imaging conditions were as follows: scan area = 200 × 100 nm^2^; number of pixels = 100 × 50; A_0_ = 2.5 nm; A_s_ = 2.1 nm. All measurements were repeated 26–51 times for each plot. Error bars represent 95% confidence intervals estimated via bootstrapping. **c,** HS-AFM images of an actin filament captured at 33 fps under feedback control with (upper) and without (lower) AGT.

### Enhancing *f_c_* by mass-control

It is extremely difficult to fabricate shorter cantilevers with *f*_c_ ≥ 3 MHz in water and *k*_c_ ≤ 0.3 N/m. To circumvent this issue, we developed a feedback control technique to enhance *f*_c_ for the spoon-shaped cantilevers. This technique is analogous to the Q-control method used for Z-scanners^27^ and cantilevers^28,29^. When the deflection signal of an oscillating cantilever characterised by the second-order transfer function *G*(*s*) is fed back into the driving signal through a transfer function *H*(*s*) (see Fig. 4a), the target second-order transfer function *T*(*s*), which connects the input and output signals, is expressed as follows:

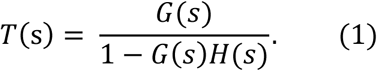

Therefore, we need to design the feedback control circuit to obtain *H*(s) characterized as follows:

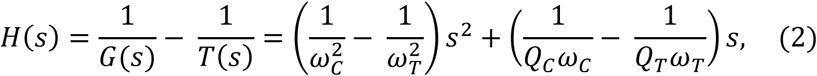

where *ω*_c_ and *ω*_T_ (*ω*_c_ < *ω*_T_) are the angular resonant frequencies, while *Q*_c_ and *Q*_T_ are the quality factors; the subscripts, C and T, denote the natural and target values of the cantilever’s mechanical properties, respectively. Eq. (2) indicates that the feedback circuit consists of the second-order and first-order derivatives in parallel. To achieve the condition of *ω*_c_ < *ω*_T_, the gain of the second-order derivative should be positive. Even when the gain of the first-order derivative is zero, *Q*_T_ can be smaller than *Q*_c_ (i.e., *Q*_T_ = *Q*_c_·*ω*_c_/*ω*_T_). Therefore, we did not use the first-derivative circuit. The addition of the second-order derivative signal to the cantilever excitation signal means that a force proportional to the acceleration of the cantilever motion is added, resulting in the reduction of cantilever’s apparent mass. Therefore, we refer to this method as ‘mass (M) control’.

**Fig. 4.**
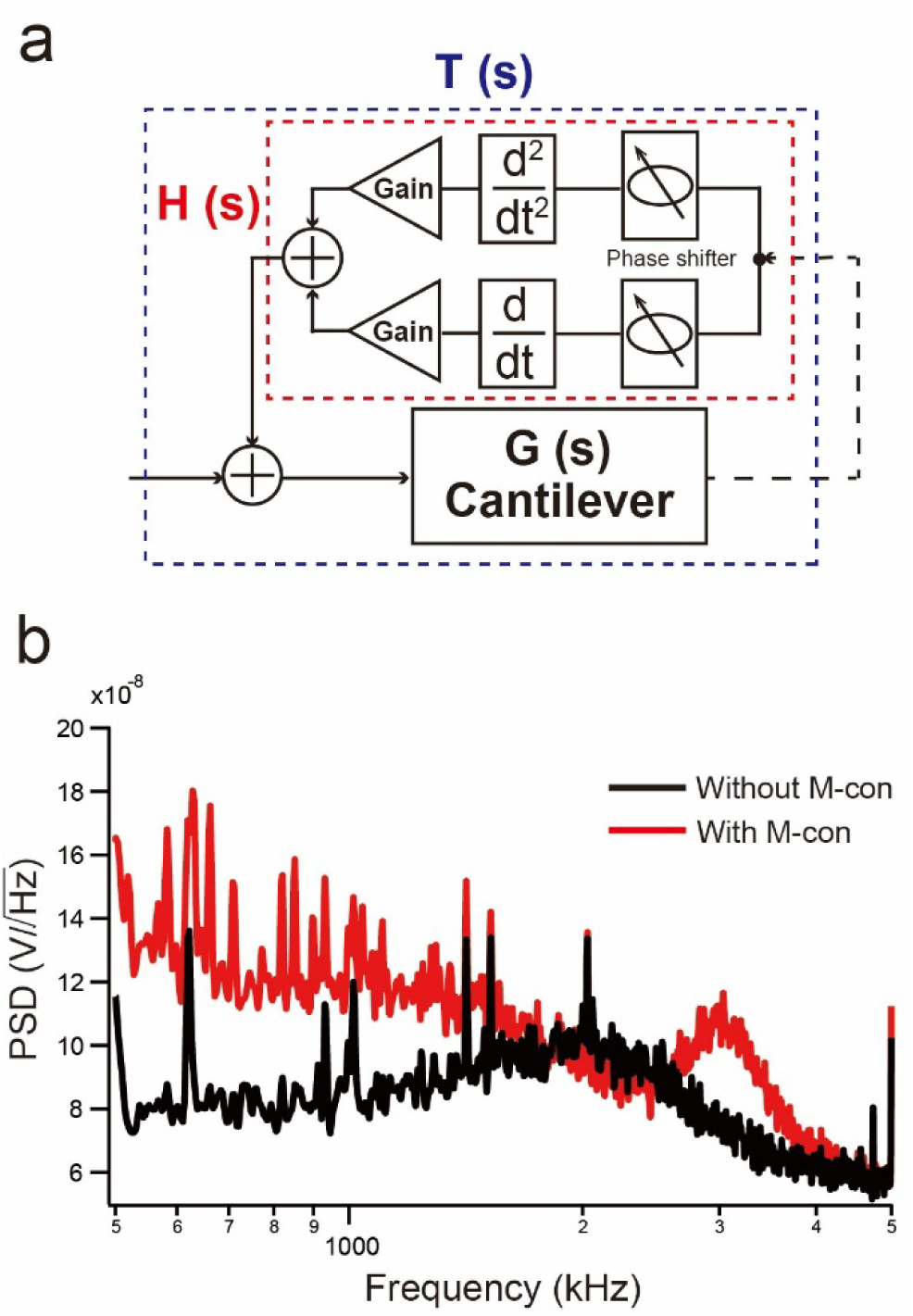
M-control for cantilevers. **a**, Schematic diagram of the M-control method. **b,** Frequency spectra of the thermal fluctuations of the spoon-shaped cantilever with (red) and without (black) M-control.

In practice, the output from the position-sensitive photodiode is differentiated using a second-derivative circuit. Its output is then fed into a phase shifter, which compensates for the phase delay inherent in the excitation of the cantilever. To minimize the phase delay, we used a photothermal cantilever excitation method^22,23^, rather than an acoustic excitation by a piezo actuator. Supplementary Figure 1 illustrates the experimental setup. The infrared laser was employed to prevent any damage to the samples^30^. To compensate for the slow response of the photothermal bending of a cantilever due to the slow heat transmission, we placed a (1 + derivative) analogue circuit corresponding to an operator of *K* × (1 + s/ω_T_) (*K*, constant gain)^27^ before the power modulation terminal of the infrared laser. The resonant frequency of the spoon-shaped cantilevers was enhanced to 3 MHz by the M-control (Fig. 4b). This level of enhancement represents the current upper limit due to the limited frequency bandwidth of the constructed analogue electric circuit.

The combination of all the above techniques has resulted in an HS-AFM system that can now image actin filaments at 33−70 fps (Fig. 5 & Movie S1), representing a significant improvement over the previous capability of 3−7 fps.

**Fig. 5.**
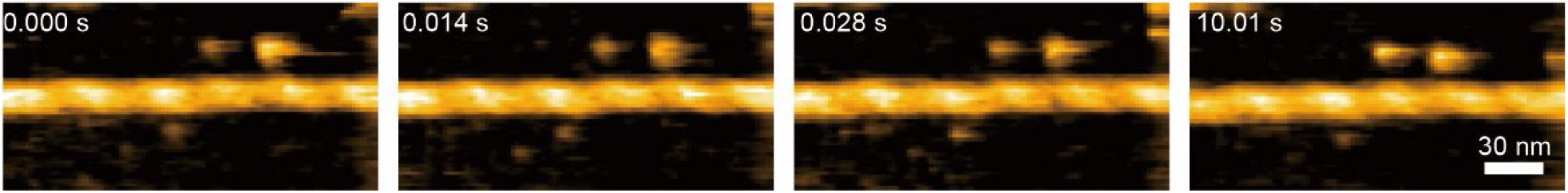
HS-AFM images of an actin filament captured at 71.4 fps. HS-AFM imaging was carried out for an actin filament oriented along the X-direction, using a combination of all techniques developed in this study. Number of pixels = 100 × 50.

### HS-AFM imaging of stator ring of F**_1_**-ATPase

The α_3_β_3_γ complex is the minimum functional unit of the rotary motor protein F_1_-ATPase^31,32^. The structural, enzymatic, and dynamic features of this complex have mostly been revealed by cryo-EM^33,34^ (or X-ray crystallography)^35–37^, biochemical analyses^38–40^, and single-molecule optical measurements of an optical marker attached to the γ-subunit^41–46^, respectively. The rotor γ-subunit is inserted into the central cavity of the hexagonal α₃β₃ stator ring, which is formed from alternately arranged α and β subunits. The catalytic sites are located at the αβ interfaces, primarily on the β-subunit. One β-subunit completes one ATPase reaction cycle in a time period of *T*_c_, during which the γ-subunit performs a full 360° rotation. The reaction at the three β-subunits exhibits a phase shift in their temporal progression, with the reaction at one β-subunit occurring one-third of *T*_c_ behind the preceding reaction at the clockwise neighbour, when viewed from the α_3_β_3_ subcomplex’s C-terminal side possessing the central cavity (hereafter, this viewing direction is commonly used). This reaction is known as ‘rotary catalysis’^47,48^. Therefore, the reactions at the three β-subunits are highly interdependent. The 120° unitary rotation of the γ-subunit is divided into 80° and 40°^43^. This means that F_1_ undergoes two-types of pause (or dwell) three times during the full 360° rotation. Determining the chemical transitions that trigger the 80° and 40° rotations is challenging due to the occurrence of multiple chemical transitions in parallel at the three β subunits. However, it is generally accepted that the 80° sub-step is triggered by ATP binding to an empty β-subunit^43^ and that the subsequent 40° sub-step is triggered by Pi release at the anticlockwise neighbour of the β containing new ATP (see Supplementary Fig. 4)^48^. The products, ADP and Pi, are released in this order^49^.

Our previous HS-AFM study revealed that the α_3_β_3_ subcomplex, lacking the γ-subunit, undergoes rotational conformational changes during the ATPase reaction cycles^50^. One β-subunit extends outwards to adopt an open (O) conformation, while the other two retract inwards to adopt a closed (C) conformation. This results in the formation of the CCO state. The position of the open β-subunit shifts anticlockwise over the three β-subunits. This finding is not in line with the γ-dictator model^51^, which states that the rotational position of the γ-subunit determines the conformation and nucleotide state of the three β-subunits. The HS-AFM study demonstrated that the stator ring alone can generate the high cooperativity among the three β-subunits for sequential power stroking, albeit with a reduced ATPase activity. The observed conformational changes in the stator ring suggest that the γ-subunit rotates by passively receiving torques from the three β-subunits. This passive rotation, as well as the rotation of shorter γ-subunits^52^, also suggests that specific interactions between amino acid side chains in the γ and β (or α) subunits are not required. This inference was confirmed through the observation that a distinct protein, bearing a shape similar to the γ-subunit, could also rotate when inserted into the stator’s central cavity^53^. Despite the valuable contribution of the HS-AFM observation to the new findings, the observation failed to detect intermediate structures between the CCO states, which would be expected from the sub-steps optically observed for the α_3_β_3_γ complex. This is likely due to an insufficient temporal resolution in the original HS-AFM system.

We imaged the α_3_β_3_ stator ring at 40 fps in the presence of ATP using the HS-AFM developed in this study (see Methods and Movies S2–S4). The ring was covalently attached to the substrate surface at the N-terminal side (see Methods). The centre of mass of each β-subunit was calculated from its X, Y, and Z coordinates in an image (see Methods). When the centres of mass of the three β-subunits obtained from many images were marked on an averaged image of the stator ring, two groups were identified: one closer to the centre of the ring and one farther away (Fig. 6a,b), corresponding to the C and O conformations, respectively. We used this separation into two groups to determine the conformation of each β-subunit. The COO and CCO states appeared alternately (Fig. 6a). Focusing on the COO state, the position of the β-subunit with the C conformation shifts anticlockwise in the next COO state. Focusing on the CCO state reveals that the position of the β-subunit with the O conformation shifts anticlockwise in the next CCO state (Fig. 6a). Although these regular transitions were mostly observed (∼70%), irregular transition patterns were also observed, such as the backward (clockwise) positional shift of the β-subunit with C or O conformations (Table S1). Hereafter, we focus on the regular transitions and use the nomenclature β^a^(q) to describe each β-subunit based on its conformation (a – C: closed, O: open) and nucleotide occupancy (q – T: ATP, D·Pi: ADP·Pi, Pi: Pi, NF: nucleotide-free, X: temporarily undetermined). To determine the nucleotide states of three β-subunits in the COO and CCO states, the alternatively proceeding reaction of COO → CCO → OCO - - - was analysed below under the restrictions that (1) the chemical states rotate counterclockwise, (2) β^C^(NF) and β^O^(T) do not exist, and (3) ADP and Pi are released in this order.

**Fig. 6.**
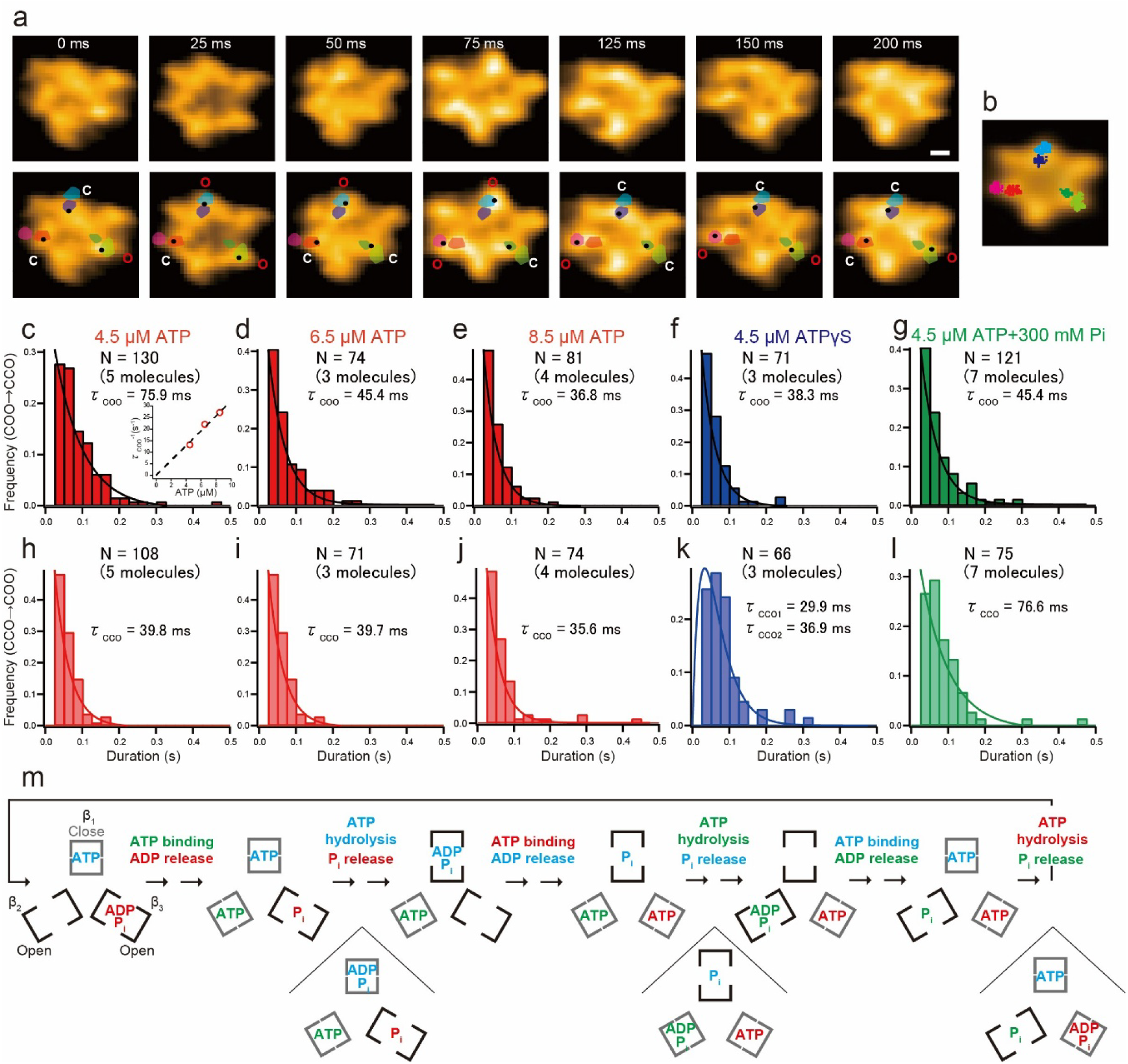
HS-AFM imaging of the α_3_β_3_ subcomplex. **a**, Successive HS-AFM images of the α_3_β_3_ subcomplex captured at 40 fps in the presence of 4.5 μM ATP. The images in upper row are shown without any labels. The centres of mass for β subunits are shown in black dots in HS-AFM images (lower row). Coloured shaded regions depicted on HS-AFM images (lower row) show the centres of mass separated into two groups corresponding to the open (O) and closed (C) conformations (see **b**). The assignment of O and C to each β subunit is denoted (lower row). Scale bar, 2 nm. **b,** The centres of mass of three β-subunits superimposed on an AFM image averaged over 200 frames are shown. Scale bar, 2 nm. **c-g,** Histograms of the COO state lifetime obtained under the indicated conditions. The hectograms were fitted to single-exponential decay functions. The fitting results are indicated in each panel. The inset in (**c**) shows an ATP-concentration dependence of the rate of transition of COO → CCO. **h-l,** Histograms of the CCO state lifetime. Solution conditions are identical to those shown in the respective upper panels (**c-g).** The histograms were fitted to single-exponential decay functions except for (**k**), while the histogram (**k**) was fitted to a double-exponential decay function assuming two consecutive first-order reactions: constant × (exp(-t/τ_1_) - exp(-t/τ_2_)). The fitting results are indicated in each panel. **m,** Scheme showing chemical and conformational transitions occurring in three β-subunits within the rotor-less F_1_-ATPase during the ATPase reaction. The three brackets represent three β subunits. Molecular species derived from the same ATP molecule are shown in the same colour.

The dwell time distributions of the COO and CCO states in the presence of 4.5−8.5 μM ATP were well fitted to single-exponential decay functions, from which their respective lifetimes (*τ*_COO_ and *τ*_CCO_) were estimated. The *τ*_COO_ lifetime depended on the ATP concentration (Fig.6c−e & inset in Fig. 6c), whereas the *τ*_CCO_ lifetime was independent. (Fig.6h−j). Therefore, β^O^(NF) exists in the COO state, but not in the CCO state. Furthermore, the CCO state following the COO state contains β^C^(T) with newly bound ATP in the same position as β^O^(NF) in the COO state.

Next, to identify the transition step at which ATP is cleaved, we imaged the stator ring in the presence of ATPγS (Movie S5). ATPγS is known to be cleaved more slowly by F_1_-ATPase than ATP^54^. At 4.5 μM ATPγS, the COO and CCO states appeared alternately (Supplementary Fig. 5). The *τ*_COO_ lifetime was not prolonged compared to the case in the presence of ATP (Fig. 6f), while the *τ*_CCO_ lifetime became significantly longer (Fig. 6k). Therefore, ATP (and ATPγS) is hydrolysed during the transition of CCO → COO. Consequently, the cleavage of ATP occurs at β^C^(T) in the clockwise neighbour of β^C^(T) with newly bound ATP, during the transition of CCO → COO. However, the dwell time distribution of the CCO state in the presence of ATPγS does not follow a single exponential function (Fig. 6k). Instead, it follows a bi-exponential function, which represents two sequential reactions. Therefore, an intermediate CCO state, which is indistinguishable in shape from its preceding CCO and contains β^C^(D·Pi), exists before transitioning into the COO state containing β^O^(D·Pi). Finally, to identify the state from which Pi is released, the stator ring was imaged in the presence of 4.5 μM ATP plus 300 mM Pi (Fig. 6g,l, Movie S6). In comparison to the 4.5 μM ATP alone case, the *τ*_COO_ lifetime was not prolonged but slightly shortened, while the *τ*_CCO_ duration was longer than that at 4.5 μM ATP alone. To check the effect of ionic strength elevated by the addition of 300 mM Pi, the stator ring was imaged in the presence of 4.5 μM ATP plus 200 mM K_2_SO_4_ (200 mM K_2_SO_4_ and 300 mM P_i_ correspond to an ionic strength of 600 and 463 mM, respectively). Despite the frequency of backward change (COO → CCO) increasing (see Table S1), no clear prolongation but slight shortening of the COO state was observed upon the addition of K_2_SO_4_ as the case of Pi addition (Supplementary Fig. 6). Therefore, it can be concluded that Pi is released during the transition of CCO → COO. Consequently, β^O^(X) solely contained in the CCO state is β^O^(Pi). Thus, all the nucleotide states of the three β-subunits in both the COO and CCO states have now been uniquely identified (see Fig. 6m). The obtained sequential reaction pattern naturally reveals that the release of ADP and Pi occurs at β^O^ but never at β^C^, and that ATP binding and ADP release, as well as ATP hydrolysis and Pi release, occur simultaneously.

The HS-AFM developed in this study revealed two conformational states (COO and CCO) transitioning during the ATPase reaction of the α_3_β_3_ subcomplex (Fig. 6m). In contrast, previous single-molecule optical measurements of the α_3_β_3_γ revealed the γ’s rotation occurring through two substeps of 80° and 40°, without any conformational information regarding the α_3_β_3_ stator ring^43,48^ (Supplementary Fig. 4). The conformational change of the β subunits drives the rotation of the γ subunit. Consequently, it can be deduced that the COO and CCO states must correspond to the substeps of the γ’s rotation. The comparison of the chemical states of the three β subunits in the two schemes (Fig. 6m and Supplementary Fig. 4), the transitions of COO → CCO and CCO → COO correspond to the substep rotations of 80° and 40°, respectively. Therefore, the chemomechanical coupling between the conformational changes in the β subunits induced by the ATPase reaction and the γ’s rotation is determined, as demonstrated in Supplementary Fig. 7.

The example of the stator ring of F_1_-ATPase demonstrates that the faster imaging attained by the method of combining techniques quantitatively reveals the complex and fast kinetics of proteins in dynamic action, which was impossible with the previous HS-AFM.

## DISCUSSION

This study presents the design and characterisation of several techniques, each of which has a moderate effect on speed performance. However, the combination of these techniques has been shown to increase the imaging rate by a factor of ∼10, at most, compared to its original instrument, while maintaining the minimised tip disturbance to the sample, spatial resolution, and the image quality. This factor of ∼10 is achieved by multiplication of the improvement factors, as 2.5 (OTI mode) × 1.4 (feedback control with ATG) × 2.0 (shorter cantilevers) × 1.5 (M-control).

The HS-AFM images of the stator ring of F_1_-ATPase acquired at 40 fps suggests potential risks when interpreting images acquired at 10 fps or less using slower apparatus. For samples whose dynamics are not as well understood as F_1_-ATPase, faster dynamics potentially occurring within the sample are not considered when interpreting the acquired images. This may lead to misinterpretation of the observed dynamics. Therefore, it is best practice to allow for a margin in the image acquisition speed (3−5 times faster capability). In addition, faster HS-AFM can expand the range of targets that can be imaged.

### Estimate of achievable improvement factors

Several designs can be developed to create the AGT capability in feedback control. However, we chose, as the first step, the simplest design in which the gain is increased when the measured amplitude *A*_m_ is lower than a threshold. This simplest design tends to generate noise in the image, particularly when the increased gain is large, limiting the speed improvement factor to 1.4. To achieve a smooth gain adjustment that is dependent on *A*_m_, a smoothly changing non-linearised error signal *ΔA*_NL_ can be created using *A*_m_ when *A*_m_ is less than the set point amplitude, *A*_s_, for example as

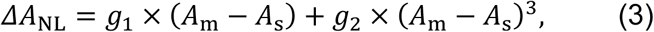

where *g*_1_ > 0 and *g*_2_ > 0 are constant gains. The non-linear error signal *ΔA*_NL_ is then fed to the conventional feedback controller. The constant gains are adjusted depending on the largest height gradient of the sample at its edge regions. This adjustment can also minimise tip parachuting. Smooth gain tuning, depending on the true error signal, *A*_m_ – *A*_s_, ensures that noise in images generated by ATG used in the simplest method is expected to be suppressed. Furthermore, the smooth gain tuning method ensures that the amplitude *A*_m_ does not decrease significantly compared to the simplest method. Consequently, the risk of sample damage can be mitigated more effectively than with the simplest method. Therefore, the smooth gain tuning method is anticipated to increase the speed improvement factor from 1.4 to at least 2.

The fabrication of AFM cantilevers shorter than ∼5 µm by chemical wet etching is extremely challenging, despite the fact that this process has been widely adopted for the large-scale production of longer cantilevers at a moderate cost. Therefore, we used 4.5 μm-long custom-made cantilevers produced using FIB milling. While this method is not suitable for mass production, the resulting products were only moderately more expensive than those produced by chemical wet etching. FIB milling is a process that allows the production of cantilevers with a length of 1 µm or less^55^. However, the OBD method of detecting cantilever deflection requires an area at the free end of the cantilever onto which a laser can be focused^2^. For this reflection area, we selected 1 × 1 µm^2^, but its mass led to a reduction of *f*_c_ in water, resulting in 2 MHz, while *k*_c_ was 0.25 N/m. It is possible to reduce the reflection area down to 0.5 × 0.5 µm^2^, which will enable *f*_c_ to be realised at 5–6 MHz in water and *k*_c_ to be maintained below 0.4 N/m. Although we need to re-design the optical system used in the OBD detector to minimise the focused laser spot, it is reasonable to expect that the improvement factor will increase from 2 to 6.

To increase the laser reflection area and resonant frequency, a design of seesaw cantilevers has recently been developed^56^. They comprise a rigid reflective board with 5 × 10 (or 5 × 5) µm^2^ oscillating over two torsional hinges with a length of 1 or 2 µm that support the board. The ample size of the reflection area is achieved with the resulting mechanical properties of 520 kHz and 0.36 N/m or 1.7 MHz and 25.6 N/m for *f*_c_ in water and *k*_c_, respectively.. Nevertheless, the seesaw cantilevers have the potential to achieve a higher *f*_c_ and a smaller *k*_c_ simultaneously by optimised design.

The M-control method is effective in increasing *f*_c_ without altering *k*_c_ or fabricating new cantilevers. We succeeded in increasing *f*_c_ by 1.5-times. This moderate increase is due to the limited frequency bandwidth of the electric circuit employed. However, more advanced electric circuits can be designed to achieve higher bandwidths^57^, thereby resulting in an enhancement factor of 2−3. However, it appears highly challenging to further increase the *f*_c_ using the M-control method once shorter cantilevers with *f*_c_ = 5−6 MHz in water are fabricated.

### Towards imaging rate of 150 fps

As discussed above, the feasible upper limit of the improvement factors can be attained are 2−3 for feedback control with AGT, 5−6 for cantilevers, and 1 for M-control when f_c_ > 5 MHz in water. Consequently, the minimum attainable improvement factor in total will become 2.5 (OTI mode) × 2 (feedback control with AGT) × 5 (cantilevers) × 1 (M-control) = 25, which is 2.5 times larger than that achieved in this study. This anticipated result corresponds to the imaging rate of 83−175 fps for relatively fragile actin filaments. The highest possible imaging rate is contingent on the spatial frequency of the sample, its fragility, and its maximum height. However, a wide range of proteins will be able to be imaged by HS-AFM at 130 fps or higher.

It is imperative that we consider the speed performance of other instrumental components, including the X- and Z-scanners, the amplitude detector and the piezo driver. However, we have already developed a method for scanning the X-scanner in a triangle wave with the vertexes being slightly rounded, at half the frequency of the X-scanner’s resonant frequency^58^. The X-scanner used in this study has a resonant frequency of ∼60 kHz, which is sufficiently high to be scanned at 20 kHz to achieve 200 fps with 100 scan lines. The Z-scanner, which can be scanned at 1 MHz, has already been achieved^18^. An amplitude detector based on the peak-hold method^4^ has already been used in this study for the 3 MHz cantilever oscillation. This amplitude detector can detect amplitude changes of a cantilever oscillating at 6 MHz. The present study used a piezo driver with a bandwidth of 1 MHz for the Z-scanner. This bandwidth is sufficient to power the Z-scanner with a resonant frequency of 1 MHz, as demonstrated^18^.

## Methods

### HS-AFM

We used a lab-built HS-AFM apparatus with modifications^4,59^. A feedback controller with AGT was used. The photothermal excitation of a cantilever was achieved by introducing an infrared laser (LuxX.HSA 980-100, Omicron-Laserage Laserprodukte GmbH, Germany) capable of modulating the laser intensity at frequencies of up to 150 MHz. An optical low-pass filter (FF01-890/SP-25, Semrock, USA) was placed in front of a position-sensitive photodiode to prevent the infrared laser from interfering with the detection of cantilever deflection. Analog electrical circuits were constructed and used for the M-control (Supplementary Fig.1). A digital-to-analog convertor board (PCI3305, Interface, Japan) and a circuit were added to the HS-AFM system for OTI mode^25^. HS-AFM imaging was carried out using a software program developed in Igor Pro (Wave Metrics Inc., USA). We used a scanner that has resonant frequencies of ∼60 kHz, ∼10 kHz and ∼350 kHz for X, Y and Z directions, respectively. Their maximum displacements at 100V are ∼850 nm, ∼3.3 μm and ∼500 nm for the X, Y and Z directions, respectively. The signal to drive the X-scanner was generated through an inverse compensation method based on the transfer function of X-scanner^60^. The cantilever deflection was converted to amplitude using a peak-hold method^4^. Cantilevers of the spoon shape, developed in this study, were used. These have a spring constant of ∼0.25 N/m, resonant frequency of ∼2 MHz in water, and quality factor of ∼2.0 in water. Since the cantilevers do not have a tip, the tip was grown to a length of ∼3 μm by electron beam deposition (EBD) using a field emission scanning electron microscope (Verios 5UC, Thermo Fisher Scientific, USA)^61^. The EBD tip was then sharpened by radiofrequency plasma etching for 30 sec at 15 W in an argon gas environment using a plasma cleaner (Tergeo Plasma Cleaner, PIE Scientific, USA). Procedures for HS-AFM imaging have been described in detail in a previous article^62^.

### Measurement of non-breaking probability

For the measurement of non-breaking probability (*P*_NB_)^25^, actin filaments oriented nearly along the Y axis or largely slanted from the X-axis were imaged for an area of 200 × 100 nm^2^ with 100 × 50 pixels at different X-scan velocities for 10 sec. We counted the number of imaging experiments that resulted in sample breakage (N_B_) and those that did not (N_NB_), and we obtained *P*_NB_ as *P*_NB_ = N_NB_ / (N_B_+ N_NB_).

### HS-AFM imaging of actin filaments

Actin was prepared from rabbit skeletal muscles as described previously^63^. Actin filaments were mixed with an equimolar amount of phalloidin (Thermo Fisher, USA). For HS-AFM imaging of actin filaments, freshly cleaved mica was treated with 0.1% 3-aminopropyl triethoxysilane (APTES, Sigma Aldrich, USA) for 3 min. After the surface was washed with pure water, actin filaments (1 μM) were deposited on the surface and incubated for 10 min. After the surface was washed with buffer A (20 mM imidazole-HCl (pH 7.6), 100 mM KCl, 2 mM MgCl_2_, and 1 mM EGTA), HS-AFM imaging was performed in buffer A.

### HS-AFM imaging of α_3_β_3_ subcomplex

The α(His_6_ at N-terminus/C193S)_3_β(His_3_-Lys_7_ at N-terminus)_3_ subcomplex of the F_1_-ATPase from thermophilic *Bacillus sp*. PS3 was expressed in *E. coli* and purified using Ni^2+^-NTA affinity chromatography and size exclusion chromatography as described previously for purification of the α_3_β_3_γ subcomplex^64^. The nucleotide-free α_3_β_3_ subcomplex was stable and stored at room temperature before use.

HS-AFM imaging of the α_3_β_3_ subcomplex was performed as described previously with minor modifications^50^. Briefly, to covalently attach the α_3_β_3_ subcomplex onto a substrate surface, we first treated the mica surface with APTES (0.05–0.1%) for 3 min and washed the surface with pure water. The mica was then treated with glutaraldehyde (0.1–0.25%) for 3 min and washed with buffer D (10 mM HEPES-NaOH (pH 7.4), 10 mM KCl, 5 mM MgSO_4_). A droplet containing the α_3_β_3_ subcomplex (1–10 nM) was deposited on the surface for 5 min, which was then washed with buffer E (10 mM Tris-HCl (pH 8.0), 2 mM MgCl_2_) to quench the reaction. HS-AFM imaging was performed in buffer F (10 mM MOPS-KOH (pH 7.0), 200 mM KCl, 2 mM MgCl_2_) under different conditions: 4.5−8.5 μM ATP; 4.5 μM ATPγS; 4.5 μM ATP plus 100−300 mM Pi; 4.5 μM ATP plus 100−200 mM K_2_SO_4_. We used a scan area of approximately 45 × 22 nm^2^ with 100 × 50 pixels. The frame rate was set to 40 fps.

### Image analysis and processing (α_3_β_3_ subcomplex)

Firstly, each target molecule was tracked using two-dimensional (2D) correlation analysis to compensate for the slow drift of the sample stage position in the lateral direction. An average filter was then applied to each tracked image to reduce noise^50^.

We assigned the conformation of the β-subunits based on the centre of mass⁶⁵. First, we drew a region of interest (ROI) enclosing the entire β subunit (approximately 10 × 10 pixels), then we calculated the centre of mass within the ROI using the X, Y and Z coordinates of the pixels contained within it. In the subsequent frame, an ROI of the same size was moved slightly to encompass the entire β-subunit, and the centre of mass was calculated. After plotting all centres of mass in the lateral plane, we applied k-means clustering criteria to divide them into two groups corresponding to open and closed conformations. Images in which the centre of the whole molecule had shifted due to drift compensation errors or image deterioration were omitted from the analysis.

## Acknowledgements

This work was supported by the JSPS KAKENHI (22H00405 to T.A. and 24K01996 to R.I.), the Human Frontier Science Program Grant (RGP0019 to T.A.), and the Shibuya Science Culture and Sports Foundation Grant to S.F.

## Author contributions

T.A. designed this project. S.F. carried out all HS-AFM experiments and image analyses.

A.O. and R.I. performed the preparation of F_1_-ATPase sample. S.F. wrote the initial draft.

T.A. wrote the final manuscript based on the discussions performed among all authors

## Competing interests

We have no competing interests.

## Additional information

Supplementary information is available for this paper.

## Supplementary Information

**Supplementary Fig. 1.**
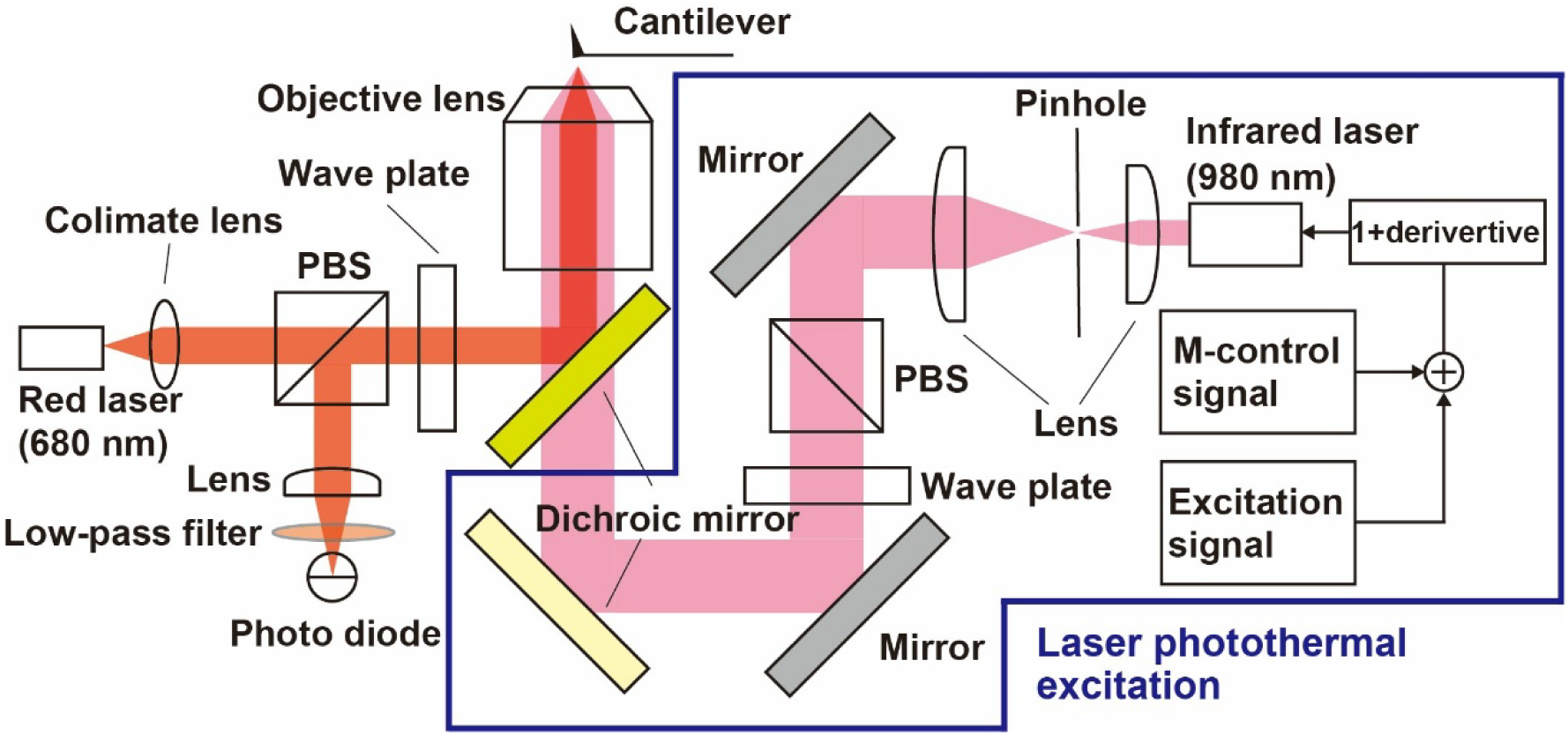
Experimental setup for the photothermal excitation of cantilever under M-control. The optics for photothermal excitation and OBD detection of cantilever deflection are shown, together with block diagrams of the electronics (M-controller and phase compensator).

**Supplementary Fig. 2.**
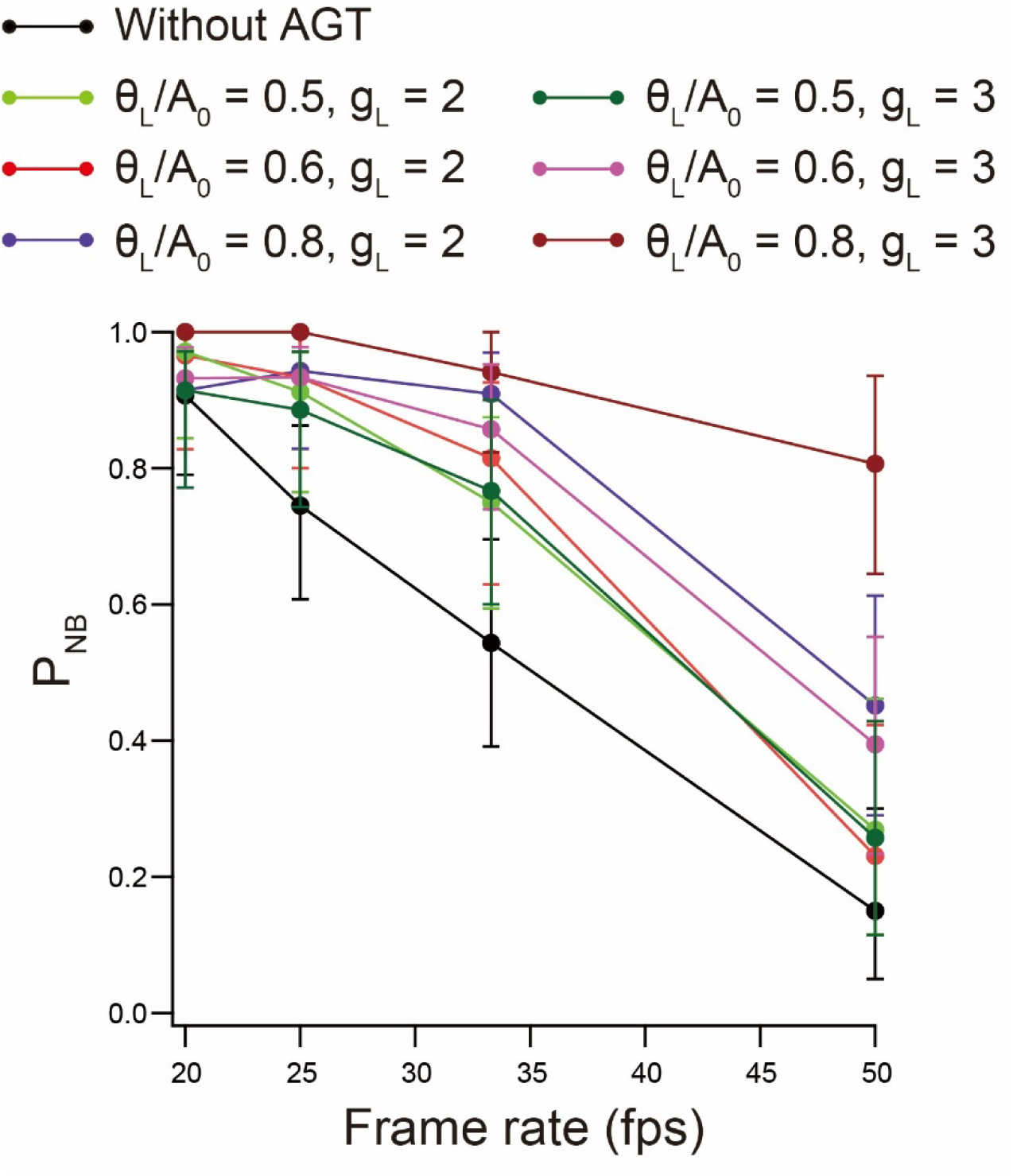
*P*_NB_ attained using AGT feedback control under various conditions. The relationship between *P*_NB_ and the imaging rate was measured by imaging actin filaments aligned along the Y-direction or slanted largely from the X-direction, using AGT feedback control with different values of *θ*_L_ and *g*_L_. The spoon-shaped cantilevers developed in this were used. All measurements were repeated 26–51 times for each plot. Error bars represent 95% confidence intervals estimated via bootstrapping. The imaging conditions were as follows: scan area = 200 × 100 nm^2^; number of pixels = 100 × 50; *A*_0_ = 2.5 nm; *A*_s_ = 2.1 nm.

**Supplementary Fig. 3.**
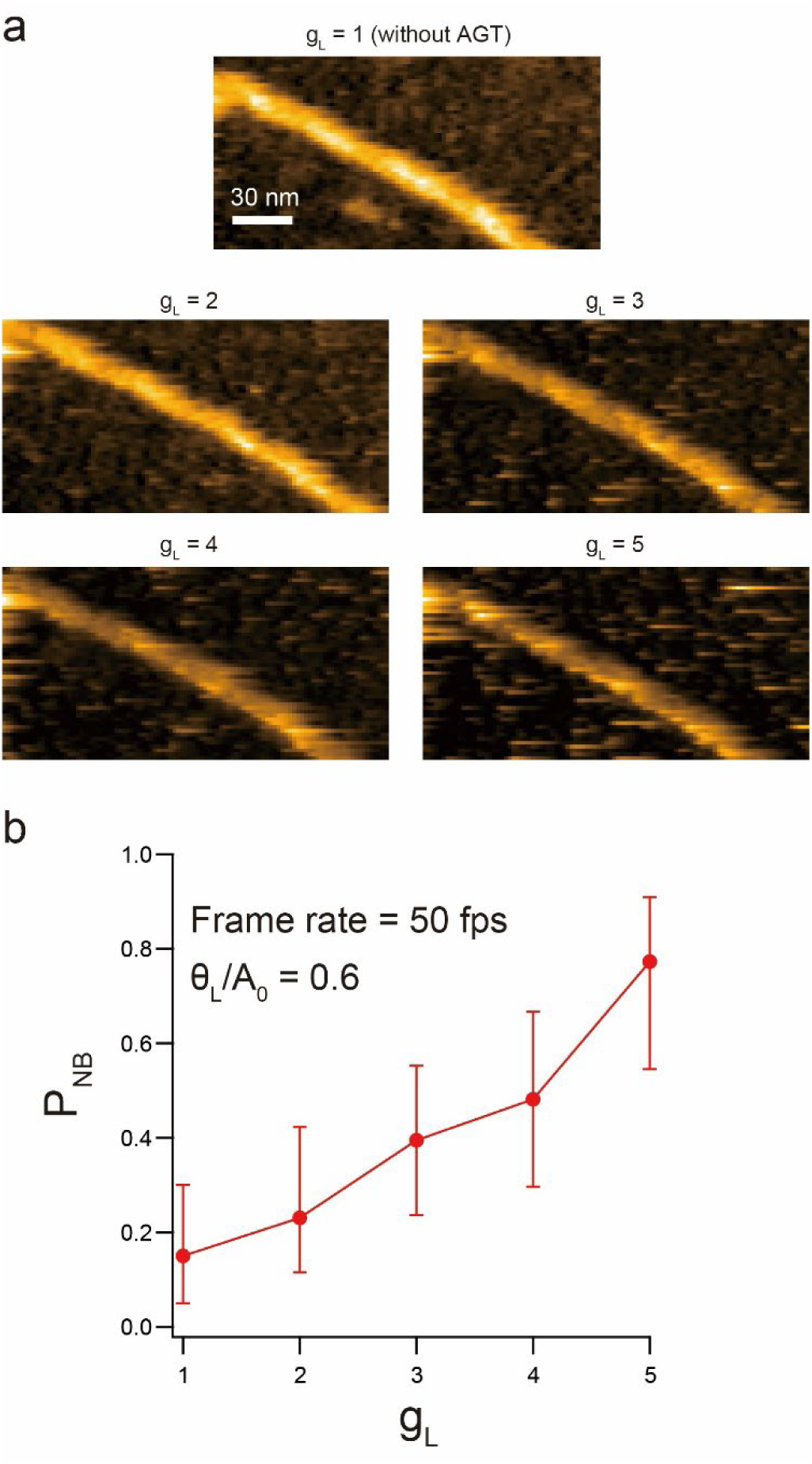
Enhancement of *P*_NB_ by increased feedback gain *g*_L_. **a**, HS-AFM images of an actin filament captured at 50 fps under AGT feedback control with variable *g*_L_. **b,** Relationships of *P*_NB_ vs *g*_L_ obtained by 50 fps imaging of actin filaments aligned along the Y-direction or largely slanted from the X-direction under AGT feedback control. All measurements were repeated 22–38 times for each plot. Error bars represent 95% confidence intervals estimated via bootstrapping. The imaging conditions were as follows: scan area = 200 nm × 100 nm; number of pixels = 100 × 50; *A*_0_ = 2.5 nm; *A*_s_ = 2.1 nm; *A*_s_/*A*_0_ = 0.8; *θ*_L_/*A*_0_ = 0.6.

**Supplementary Fig. 4.**
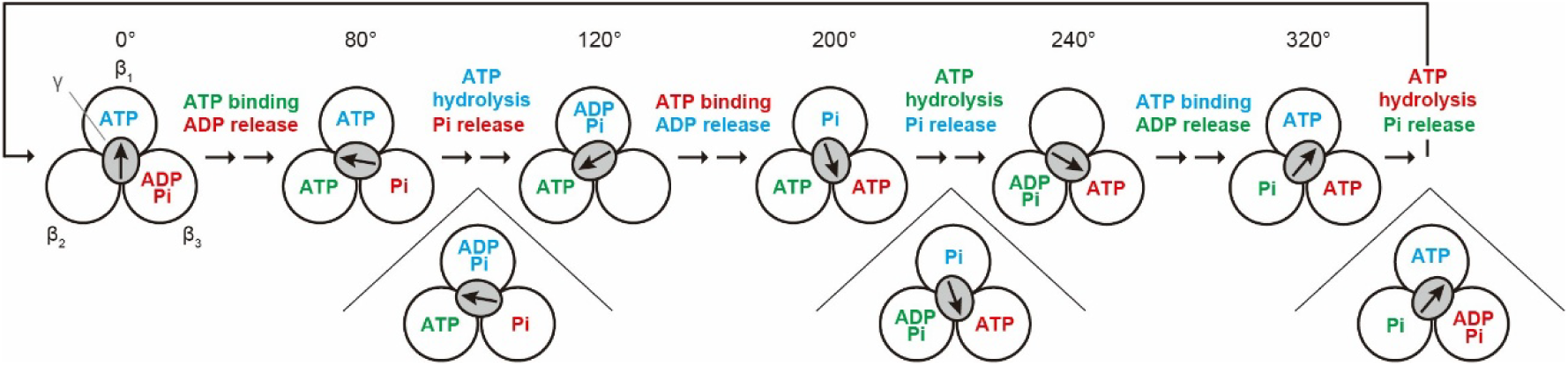
Chemomechanical coupling between γ’s rotation and ATPase reaction of F_1_-ATPase (α_3_β_3_γ). A well accepted coupling scheme of F_1_-ATPase deduced from single-molecule optical measurements is shown. The three circles represent three β subunits, while the central gray ellipsoid represents the γ subunit. The arrows in the ellipsoid show the γ’s orientation. Molecular species derived from the same ATP molecule are shown in the same color.

**Supplementary Fig. 5.**
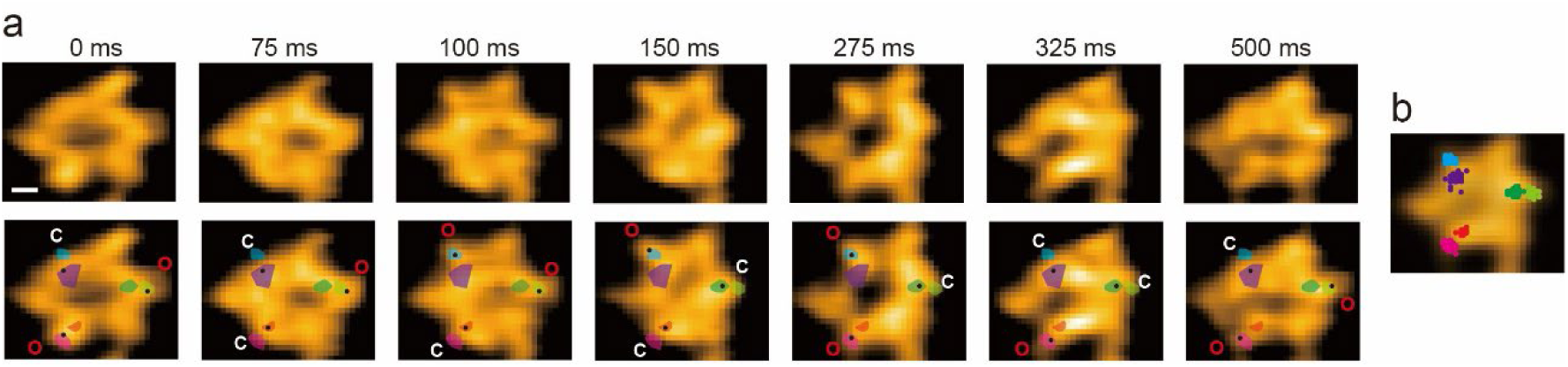
Successive HS-AFM images of the α_3_β_3_ subcomplex in the presence ATPγS. **a**, The images were captured at 40 fps in the presence of 4.5 μM ATPγS. The centres of mass of β subunits are shown in black dots on the HS-AFM images (lower). Coloured shaded regions in lower panels show that the centres of mass are separated into two groups corresponding to the open (O) and closed (C) conformations (see **b**). The O and C conformations of β-subunits are denoted in HS-AFM images (lower panels). Scale bar, 2 nm. **b,** The centres of mass of β-subunits superimposed on an AFM image obtained by averaging 100 frames are shown.

**Supplementary Fig. 6.**
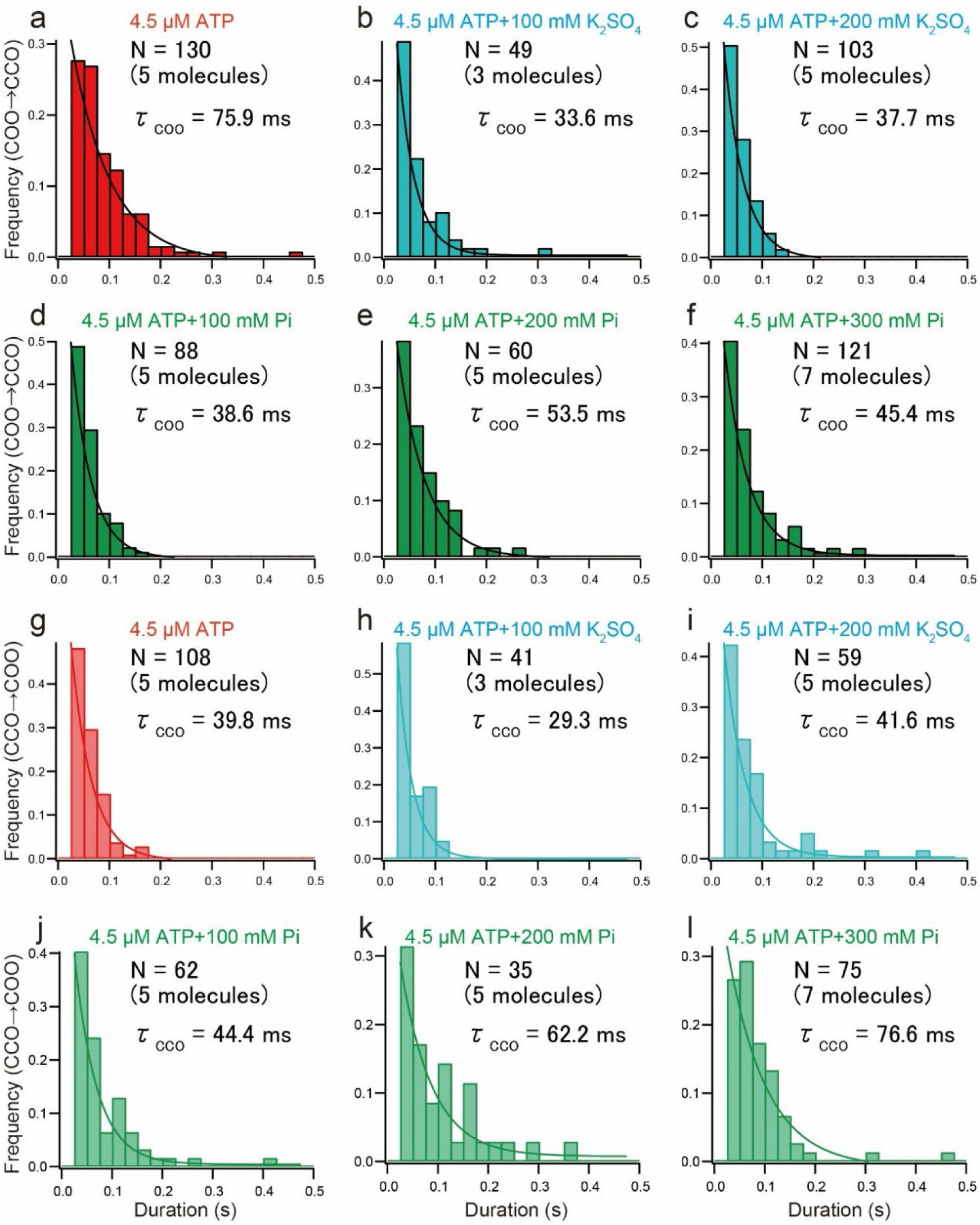
Effects of ionic strength on the conformational transitions of the α_3_β_3_ subcomplex observed under different conditions. All imaging experiments were carried out at 4.5 μM ATP. The concentrations of Pi and K_2_SO_4_ were indicated in the respective panels. **a-f,** Dwell time distributions for the COO state. The distributions were fitted to single-exponential decay functions. **g-l,** Dwell time distributions for the CCO state. The distributions were fitted to single-exponential decay functions.

**Supplementary Fig. 7.**
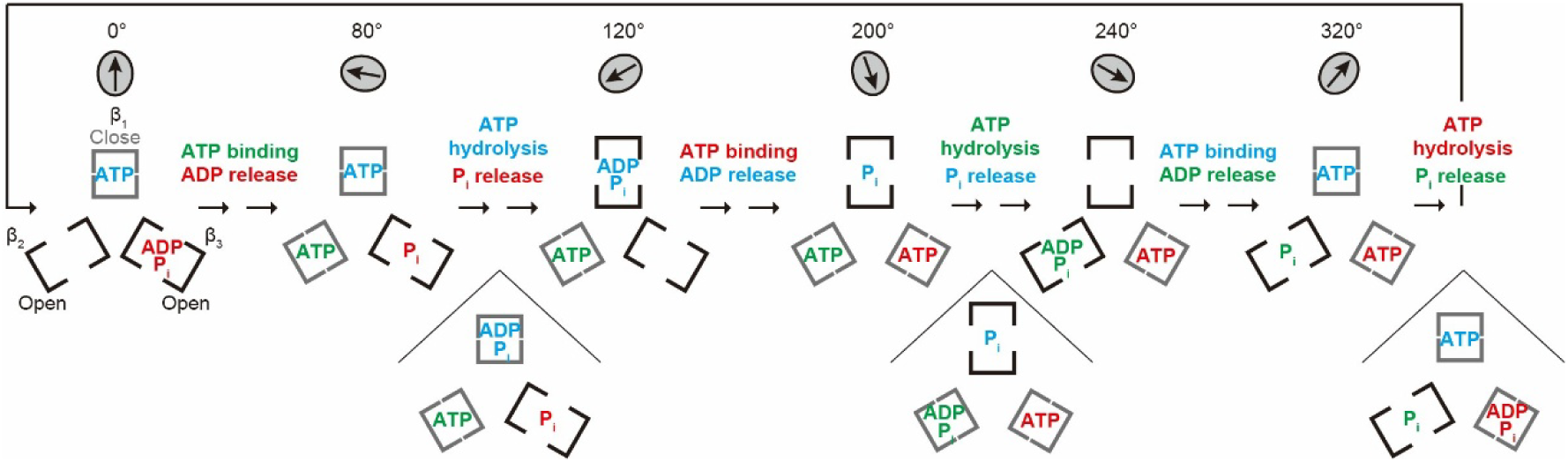
Relationship between the βs’ conformational transitions in the α_3_β_3_ subcomplex and the substep rotations of γ-subunit in the α_3_β_3_γ complex during ATPase reaction. The relationship between the β’s conformational states and the chemical states shown in Fig. 4 is used. The relationship between the γ’s substep rotations and the chemical states is obtained from previous single-molecule optical measurements. The three brackets represent three β subunits with open or closed conformations. Molecular species derived from the same ATP molecule are shown in the same colour. Angle changes of the γ subunit within the α_3_β_3_γ complex are shown on the top.

**Table S1.** Conformational changes of the α_3_β_3_ subcomplex observed in this study.

|  | Forward<br>COO→CCO<br>(%) | Forward<br>CCO→COO<br>(%) | Backward<br>CCO←COO<br>(%) | Backward<br>COO←CCO<br>(%) | Forward<br>COO→COO<br>(%) | Forward<br>CCO→CCO<br>(%) | Backward<br>COO←COO<br>(%) | Backward<br>CCO←CCO<br>(%) | Total<br>Number<br>(N) |
| --- | --- | --- | --- | --- | --- | --- | --- | --- | --- |
| 4.5 μM ATP | 37.2 | 31.8 | 6.0 | 10.3 | 8.6 | 4.0 | 1.1 | 0.9 | 349<br>(5 molecules) |
| 6.5 μM ATP | 29.7 | 28.5 | 14.5 | 10.4 | 8.4 | 6.0 | 1.2 | 1.2 | 249<br>(3 molecules) |
| 8.5 μM ATP | 32.3 | 29.9 | 6.8 | 6.4 | 13.1 | 6.0 | 2.8 | 2.8 | 251<br>(4 molecules) |
| 4.5 μM ATPγS | 31.0 | 28.4 | 12.1 | 12.9 | 2.6 | 8.6 | 1.7 | 2.6 | 232<br>(3 molecules) |
| 4.5 μM ATP<br>+ 100 mM P<sub>i</sub> | 39.6 | 27.9 | 6.8 | 12.2 | 6.3 | 4.5 | 1.8 | 0.9 | 222<br>(5 molecules) |
| 4.5 μM ATP<br>+ 200 mM P<sub>i</sub> | 41.3 | 21.3 | 5.8 | 21.3 | 4.5 | 5.2 | 0.6 | 0 | 155<br>(5 molecules) |
| 4.5 μM ATP<br>+ 300 mM P<sub>i</sub> | 41.3 | 25.2 | 4.4 | 17.4 | 3.0 | 5.0 | 2.7 | 1.0 | 298<br>(7 molecules) |
| 4.5 μM ATP<br>+ 100 mM K<sub>2</sub>SO<sub>4</sub> | 29.9 | 25.6 | 7.9 | 14.0 | 6.7 | 10.4 | 1.8 | 3.7 | 164<br>(3 molecules) |
| 4.5 μM ATP<br>+ 200 mM K<sub>2</sub>SO<sub>4</sub> | 38.7 | 23.6 | 4.4 | 18.5 | 5.9 | 6.6 | 0.3 | 1.8 | 271<br>(5 molecules) |

Others conformational changes involving the CCC, OOO and shifts that skip the two elementary steps were omitted in the analysis.

## Description of Additional Supplementary Files

**Supplementary Movie S1 |** HS-AFM movie of an actin filament oriented along the X-direction, recorded at 71.4 fps. The movie was captured for a scan area of 200 × 100 nm^2^ with 100 × 50 pixels.

**Supplementary Movie S2 |** HS-AFM movie of the α_3_β_3_ subcomplex in the presence of 4.5 μM M ATP, recorded at 40 fps. The movie was captured for a scan area of 45 × 22.5 nm^2^ with 100 × 50 pixels, and cropped to 17× 15 nm^2^.

**Supplementary Movie S3 |** HS-AFM movie of the α_3_β_3_ subcomplex in the presence of 6.5 μM ATP, recorded at 40 fps. The movie was captured for an scan area of 45 × 22.5 nm^2^ with 100 × 50 pixels, and cropped to 17× 15 nm^2^.

**Supplementary Movie S4 |** HS-AFM movie of the α_3_β_3_ subcomplex in the presence of 8.5 μM ATP, recorded at 40 fps. The movie was captured for a scan area of 45 × 22.5 nm^2^ with 100 × 50 pixels, and cropped to 17× 15 nm^2^.

**Supplementary Movie S5 |** HS-AFM movie of the α_3_β_3_ subcomplex in the presence of 4.5 μM ATPγS, recorded at 40 fps. The movie was captured for a scan area of 45 × 22.5 nm^2^ with 100 × 50 pixels, and cropped to 14× 12 nm^2^.

**Supplementary Movie S6 |** HS-AFM movie of the α_3_β_3_ subcomplex in the presence of 4.5 μM ATP plus 300 mM P_i_, recorded at 40 fps. The movie was captured for a scan area of 45 × 22.5 nm^2^ with 100 × 50 pixels, and cropped to 16× 15 nm^2^.

## Notes

### Competing Interest Statement

The authors have declared no competing interest.

